# Specific cholesterol binding drives drastic structural alterations in Apolipoprotein A1

**DOI:** 10.1101/228627

**Authors:** Arjun Ray, Asmita Ghosh, Rahul Chakraborty, Santosh Kumar Upadhyay, Souvik Maiti, Shantanu Sengupta, Lipi Thukral

## Abstract

Proteins typically adopt a multitude of flexible and rapidly interconverting conformers, many of which are governed by specific protein interaction domains. While disc-shaped oligomeric HDL and its major protein component ApoA1 have been the focus of several investigations, yet the structural properties of monomeric ApoA1 remain poorly understood. Using tens of independent molecular simulations (>50 μs), we reveal that ApoA1 adopts a compact conformation. Upon addition of a physiological concentration of cholesterol to ApoA1, the monomeric protein spontaneously formed a circular conformation. Remarkably, these drastic structural perturbations are driven by a specific cholesterol binding site at the C-terminal and a novel cholesterol binding site at the N-terminal. We propose a mechanism whereby ApoA1 opens in a stagewise manner and mutating the N-terminal binding site destroys the open ‘belt-shaped’ topology. Complementary experiments confirm that the structural changes are induced by specific association of cholesterol with ApoA1, not by the non-specific hydrophobic effect.

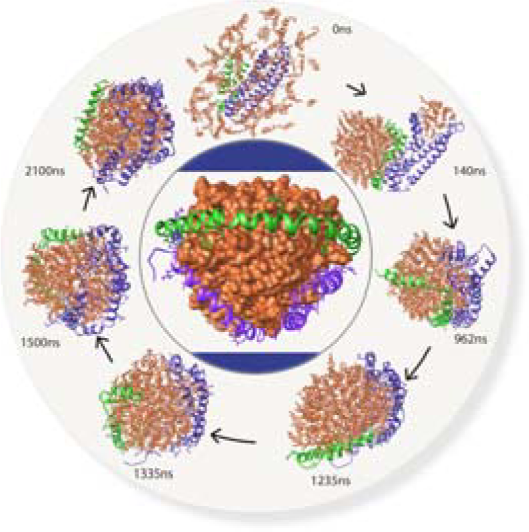
Toc Graphics.

---

The removal of cholesterol in humans is universally operated by reverse cholesterol pathway (RCT), which tightly controls lipid homeostasis via the transfer of cholesterol from peripheral cells to the liver^1-3^. A pivotal step in the formation of high-density lipoprotein (HDL) complex involves sequestering of cholesterol by apolipoproteins. Apolipoprotein A1 also commonly referred to as ApoA1, is the major protein constituent of HDL (~70%)^4^. Although a relatively small protein of 243 amino acids, ApoA1 exhibits remarkable structural variability with varying degrees of cholesterol associated protein states^5^. These distinct structural states, range from lipid-free monomeric conformation in blood plasma cells^6^, low-concentration cholesterol-bound monomeric conformation^7-8^, to higher oligomeric forms involved in belt-shaped HDL supramolecular assemblies^9-12^. Given the association of elevated cholesterol with chronic cardiovascular diseases, understanding the structural basis of ApoA1 functional diversity is a long-standing open question in the field.

Although direct crystallographic evidence is lacking for monomeric ApoA1 protein, there exist several reports highlighting the aspects of ApoA1 structure and function. The ApoA1 structure, suggested by proteolysis and mutagenesis studies is divided into two structural domains i.e., N- and C-terminal that are highly conserved across species (Figure S1). A recent hydrogen exchange (HDX) study suggested highly helical N-terminal domain and disordered residues within C-terminal^13^. Further the degree of total helical content of the protein in apo and bound forms has been shown to vary using several biophysical approaches^15-16^. A putative close proximity of interaction between the N- and C-terminal is also proposed^14^. Recently, a chimeric ApoA1 monomeric structure has been proposed based on a consensus approach^17^. While these studies have characterized many aspects of monomeric structure, a conclusive model and direct structural evidence on interconverting lipid-free and cholesterol-bound states is largely missing.

This study is aimed at investigating the structural properties of ApoA1 upon its interaction with cholesterol. This is undoubtedly the most important step in biogenesis of HDL formation. Our approach was twofold. We probed ApoA1 protein structural variations with and without cholesterol in all atomistic representation and, next highlighted observed dynamic differences within protein domain elements that may be intricately coupled with its biological function. First, to understand the functional significance of lipid-free ApoA1, we generated the ApoA1 starting structure using a hybrid modeling and simulation protocol (see Methods and Figure S2). Further, we performed multiple atomistic molecular dynamics (MD) simulations (5x7 μs), leading to determination of structure and dynamics with statistical reliability. The mapped structural dynamics at μs timescale is three order magnitude longer than the previous reports^18-19^.

To fully describe the conformational changes, we utilized angle calculation between N- and C-terminal domains as an order parameter (Figure 1A). As a function of time, the two domains reached proximity from ~100° to 40° within 5 μs, and remained stable for the last 2 μs in all simulations. Remarkably, in each of the five simulations, ApoA1 converged to a closed V-shaped topology (Figure S3). Figure 1B shows the time evolution towards the final structure, with intact N- and C-terminal domains. In addition, structural properties of V-shaped topology ensemble were analyzed and we found slight intra-domain bending variations (Figure S4). The predicted dynamic properties are in good agreement with previously reported site-directed electron paramagnetic resonance (EPR) spectroscopy^20^, where the probes localised at residues 163, 217, and 226 revealed close association and parallel topology between N- and C-terminal (Figure 1C). This structure obtained from extensive MD simulations thus enabled us to study protein-cholesterol interaction with high reliability.

**Figure 1:**
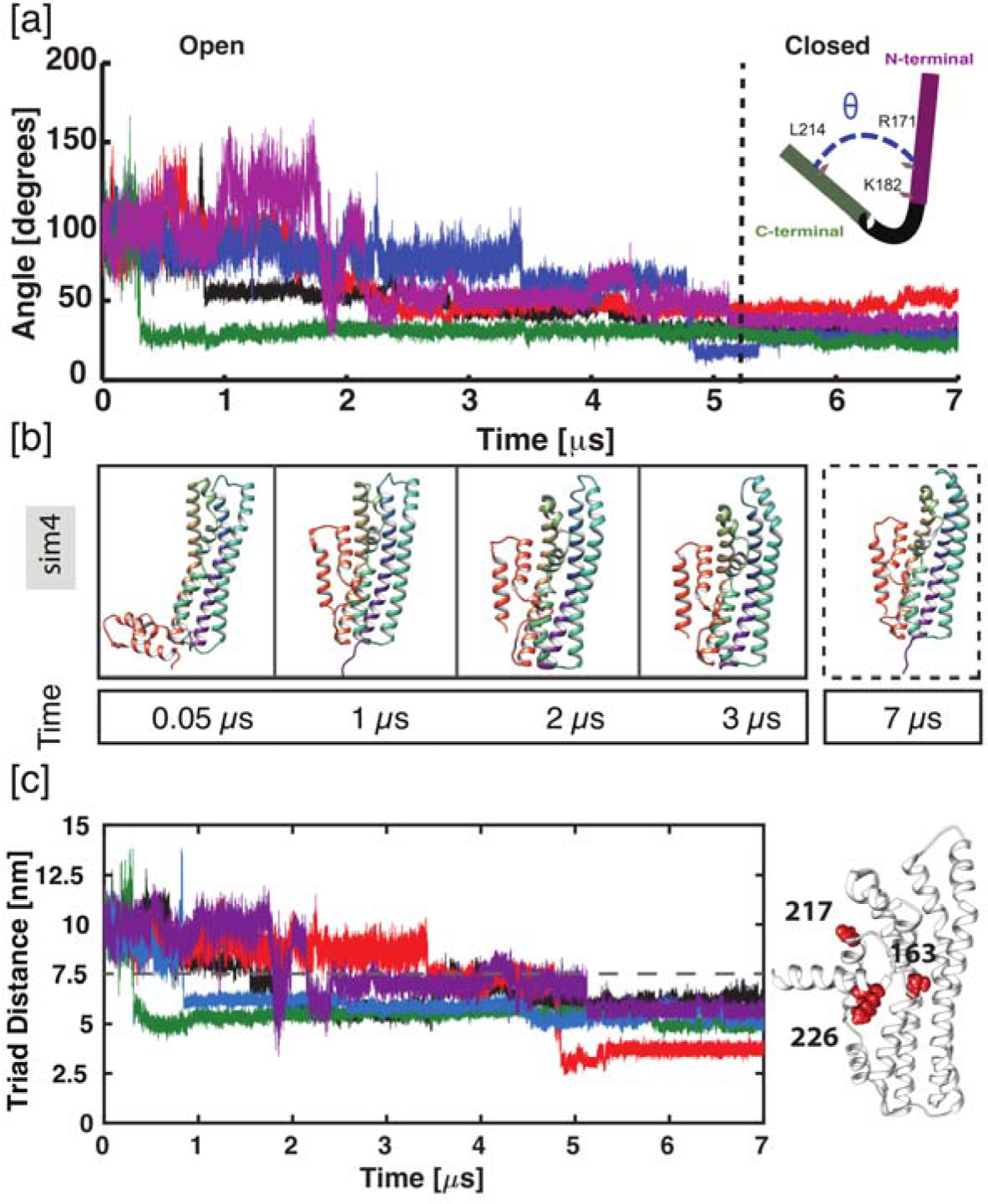
ApoA1 Lipid-free Structure and Dynamics: A) Time evolution of ApoA1 structure as a function of angle between C- and N-terminal domains. The angle was calculated based on three residues (171, 182 and 214) across all simulations, as highlighted in the schematic diagram. B) Snapshots of ApoA1 structures at various time points during one of the representative simulation showed a V-shaped topology towards the end of simulation time. The colour scale represents different helices. C) In accord with the experimental data reported^20^, time evolution of the triad distance is converged to <=7.5 nm, which represents V-shaped topology. The location of the three residues is marked on the ApoA1 structure.

It is well characterized that ApoA1 exhibits protein regions that are likely to be essential for its cholesterol-associated physiological function. At the atomic level, however, our understanding is largely obscure due to fast-timescale conformational transitions that are experimentally difficult to characterize. Therefore, we assessed how cholesterol guides structural changes in monomeric ApoA1 protein. The evolution of dynamic structural changes with residue-based fluctuations are shown in Figure 2A. There are two major observations. First, cholesterol binds protein at two distinct sites that are subsequently merged at ~400 ns, followed by opening up of ApoA1 structure to fully sequester the cholesterol. Second, the domain-wise protein mobility is highly variable. While the N-terminal is stable and well defined, the C-terminal region is highly mobile. At 1315 ns, the C-terminal twists and forms two short stretches of helix, before opening up and forming a circular or open topology. By the end of the simulation length, we found that all the residues interact with lipids, with the cholesterol in the center. These findings suggest that ApoA1 dynamically modulates its topology in the presence of cholesterol.

**Figure 2:**
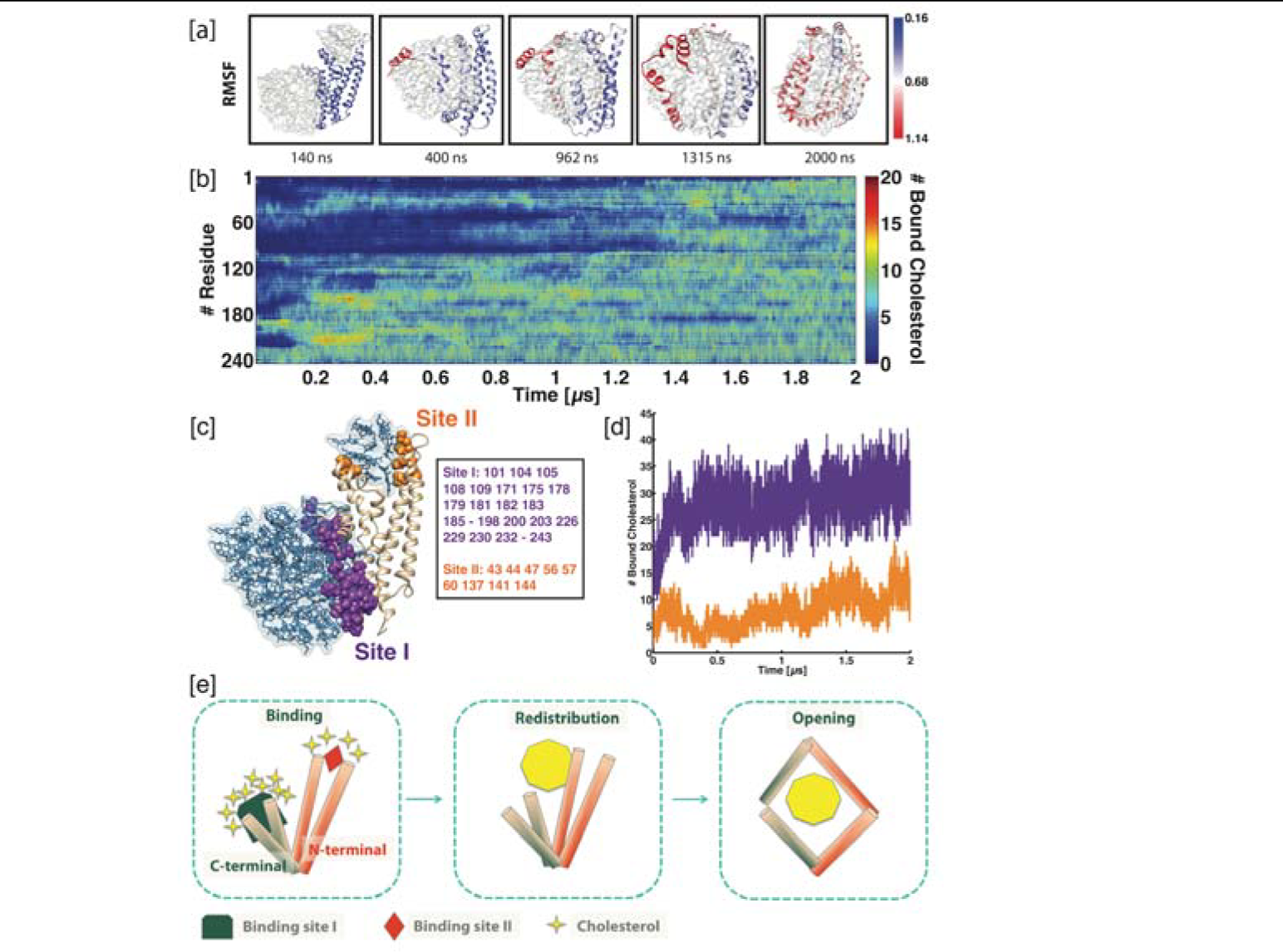
Dual topological organization of ApoA1 upon cholesterol binding. A) The representative snapshots are shown as a function of time for ApoA1-cholesterol simulations. The residue-based fluctuations (RMSF) are mapped onto the structure and the corresponding colour bar denotes the RMSF values in nm. B) The cholesterol distribution on each protein residue is shown along the entire simulation length. C) Representation of two major cholesterol binding sites and the participating residues. Site I refers to the binding site encompassing C-terminal while Site II refers to the binding site at N-terminal, shown in purple and orange, respectively. D) Time evolution of the two major cholesterol binding sites showed specific initial binding. E) Collectively, the events during ApoA1 cholesterol association can be divided into three stages, as shown in schematic representation.

To quantitatively assess the interaction, we plotted the time occurrence of the cholesterol interaction with ApoA1 (Figure 2B). During the initial simulation time (140 ns), cholesterol was found to specifically target two binding sites. The C-terminal is the main site of cholesterol targeting while the second major binding pocket is at the top of the N-terminal helix bundle (Figure 2C). These two sites referred from here as Site I and Site II, respectively thus defines cholesterol binding sites with high propensity. Figure 2D shows time evolution of these two major cholesterol binding sites. There is a concomitant lipid binding at both sites followed by cholesterol movement from both terminal sites merging towards the core. Although the importance of C-terminal binding of cholesterol has been reported^21-22^, the role of N-terminal binding has not been established. To further prove the role of N-terminal binding, the hydrophobic residues at Site II were mutated to Alanine residues. The mutated simulation showed that the cholesterol does not bind at the N-terminal site, thereby limiting the mobility of C-terminal and does not lead to N-terminal opening (see SI Figure S5). This study provides further evidence that N-terminal binding is critical for selectively recognizing cholesterol.

The results described above suggested that ApoA1 cholesterol association can be divided into following stages: a) Specific cholesterol binding at two sites, b) Redistribution with cholesterol merging from two sites, and c) Complete sequestering that involves N-terminal opening along with C-terminal plasticity (Figure 2E).

We reasoned that the binding specificity of cholesterol followed by N-terminal opening could be exploited to reveal quantitative aspects of stable elements. To assure maximum sampling available to the system, we leveraged above mentioned ApoA1-cholesterol simulation for selecting six conformations from various time points of the simulation. Each of these six conformations served as starting structure for further simulation (up to 250 ns each). Six replicates of each were simulated to obtain a sizeable pool of conformations (Figure 3A). The structural characteristics of the starting structures are described in Figure S6.

**Figure 3:**
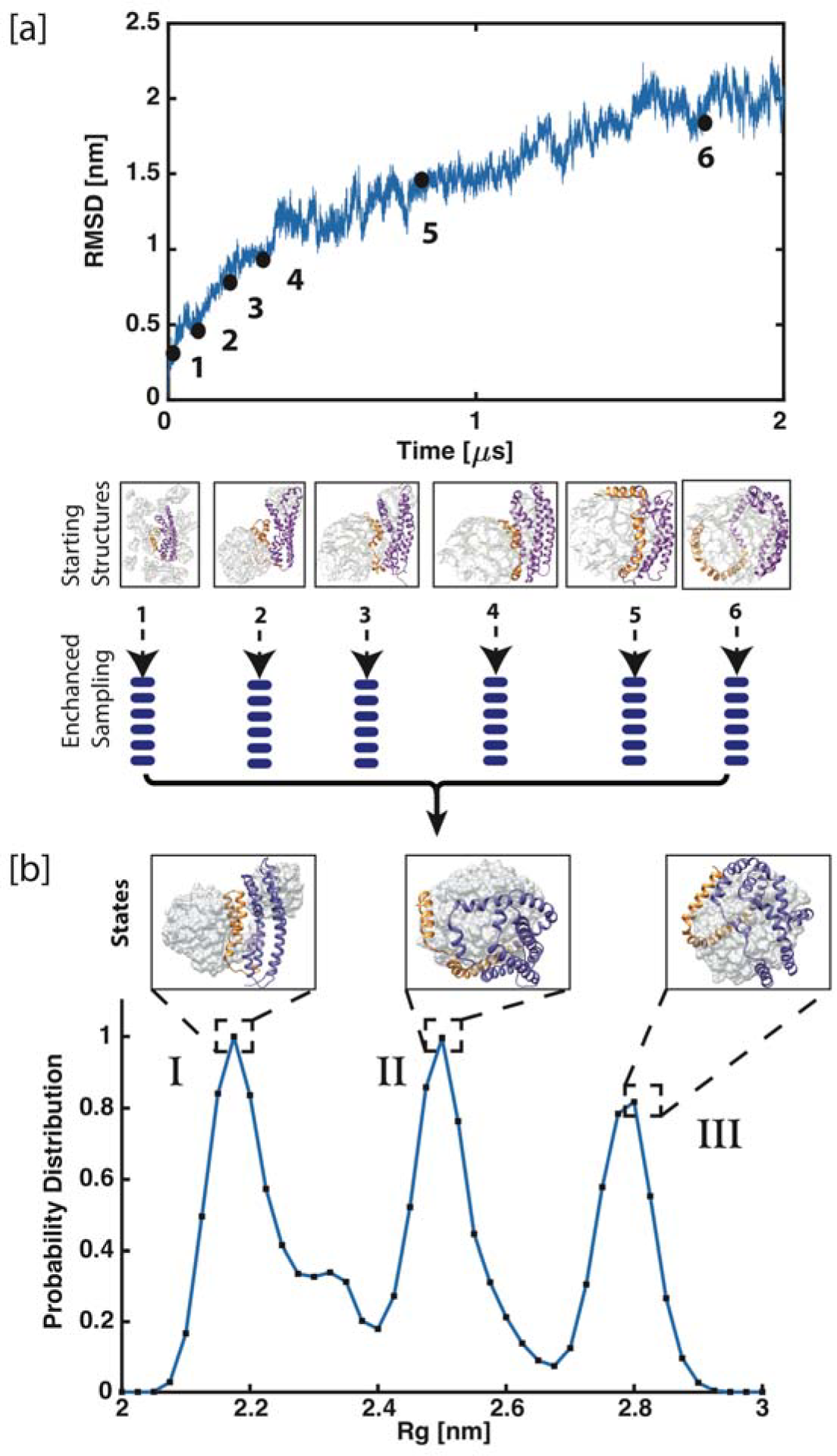
Mapping conformational space of ApoAl during closed to open structural transition. A) The time evolution of RMSD as a function of initial pilot simulation was used to extract six seed structures (highlighted in boxes). The RMSD values are calculated with reference to the initial V-shaped topology structure. In total, six replicas were started from each seed structures, to obtain increased sampling. B) The one-dimensional probability distribution is plotted as a function of radius of gyration (Rg). See methods for details. Clearly, three major states (State I-III) are populated in the map. The represented structures of each state are shown in protein ribbon diagram with N- and C-terminal colored as purple and orange, respectively. The cholesterol is marked in surface representation.

We found that there is a clear distribution of population density, as seen in the one-dimensional probability distribution graph, plotted as a function of Rg (Figure 3B). Three states (I, II and III) predominantly populate the conformational landscape, also using different order parameters (Methods; Figure S7). To quantify the population distribution, we determined the properties of structural ensemble in each of the three states. Clearly, state I and III represent closed and open topology, respectively. Intriguingly, state II was found to be one of the major cluster and multiple inter-conversion of states, 1 ↔ 2, 2 ↔ 3 were observed in our simulations (Figure S8). During transition from state I to III, a common state II was observed in all instances. In addition, there were multiple fast- and slow-transitions between respective states: 41% (I→II), 40% (I←II), 6% (II→III) and 11% (II←III). However, we never observed I↔III transition, indicating that State II is an obligatory “intermediate state” in reaching the final open conformation.

To validate the above findings observed from simulations, we performed quantitative experiments to understand ApoA1-cholesterol ligand induced binding changes. Purification of monomeric lipid-free ApoA1 poses experimental challenges due to its inherent nature of aggregation in moderately high concentration. We purified and confirmed the monomeric full-length lipid-free ApoA1 by Matrix Assisted Laser Desorption/Ionization (MALDI), with absence of any dimeric or oligomeric species (Figure S9). Dynamic light scattering (DLS) with 1 mg/ml of the protein showed significant loss of monomeric species while there was an abundant amount of dimeric and other oligomeric species that formed upon concentrating the protein. Therefore, we ensured that the protein was kept at substantially low concentration, to confirm the monomeric state of the protein in all our experiments.

In order to probe binding energetics of ApoA1 and cholesterol binding, we performed Isothermal Titration Calorimetry (ITC) at the room temperature. As shown in Figure 4A, a titration curve of cholesterol in buffer (in red) depicts a large exothermic heat signal, which gradually decreases until the 20th injection. Interestingly, the heat signals of the first few injections (7 to 8) displayed less exothermic heat in the titration of cholesterol into ApoA1 (in black) and magnitude of the heat signals become close to the titration of cholesterol into buffer at 9th injection. The integrated enthalpic curves obtained from ITC titrations, shown in Figure 4B, contain observed enthalpy changes for each injection of 0.1 mM cholesterol into buffer.

**Figure 4:**
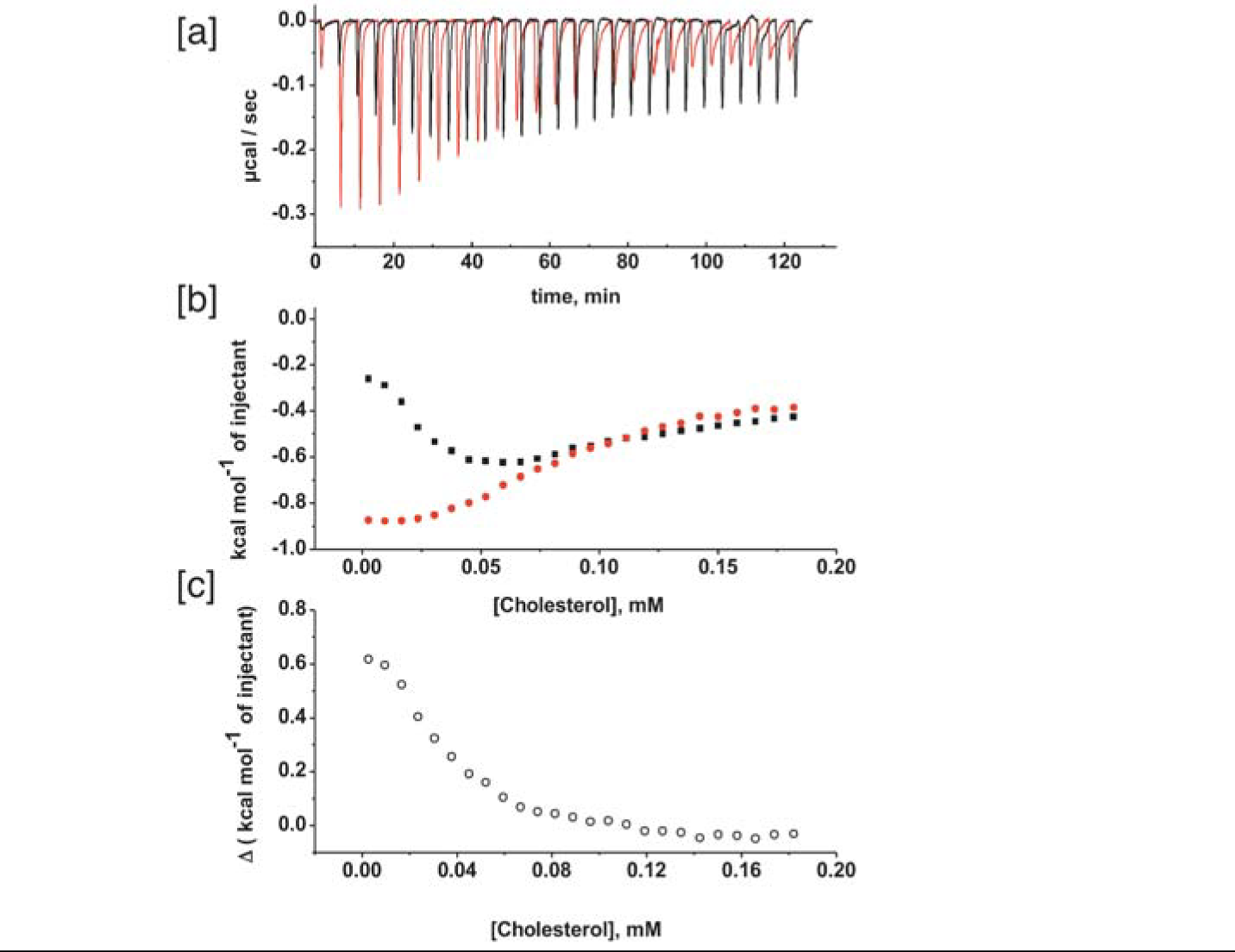
Isothermal titration calorimetry (ITC) A) The thermograms (cell feedback (CFB) in fical/s) for titrations of 1mM cholesterol into buffer (red) and 10 μM ApoA1 (black). B) The ΔH_cholesterol to ApoA1_ (black) and ΔH_cholesterol to buffer_ (red) was obtained by integrating the area under the CFB curve at each injection normalized to the amount of cholesterol injected. C) Enthalpic change due to the monomeric cholesterol association to ApoA1 as a function of cholesterol concentration.

The dilution curve of cholesterol into buffer can be classified into two regions. In region I, the injectant containing concentrated cholesterol produced large exothermic heat effects due to the deaggregation process of the aggregated cholesterol (monodispersed form). However, the integrated enthalpic curves of cholesterol titrating into ApoA1 follow a different trend. In this region, the endothermic binding of monodispersed cholesterol to ApoA1 was observed. In region II, lower exothermic contribution in each subsequent injection is indicative of relatively less deaggregation of cholesterol. However, under equilibrium monodispersed cholesterol continues to bind with unoccupied ApoA1 binding sites.

Further, we plotted the differential enthalpic values under each injection (ΔH_cholesterol to ApoA1_ ΔH_cholesterol to buffer_) as a function of cholesterol concentration (Figure 4C). As observed, enthalpy change for such interaction is endothermic in nature, and the interaction is completely entropy driven. This is required to initiate the hydrophobic interaction due to large amount of dehydration originating from protein and cholesterol binding. These findings from ITC experiments suggest that ApoA1-cholesterol interaction exhibits ligand-induced equilibrium shift between different conformational states.

We also performed an additional control experiments to probe the effect of hydrophobic driven non-specific interactions. The hydrophobic amino acid tryptophan (TRP) was selected for this purpose and the titration of ApoA1 (10 iμl, cell) and TRP (1 mM, in syringe), as shown in Figure S10. We noted that the curve revealed low heat profile, indicating negligible interaction with ApoA1. In addition, we studied the tryptophan-ApoA1 interaction at atomistic level using MD simulations. In comparison to ApoA1-CHL simulations, TRP showed minimal binding at Site 1 and 2. This data also confirmed that binding of ApoA1 and cholesterol is not a general hydrophobic effect.

We then attempted to evaluate the changes in secondary structure of ApoA1 upon cholesterol addition and how secondary structure preferences of different segments of ApoA1 vary dynamically in lipid-free and cholesterol-associated simulations. In order to probe these differences, we computed the probability distribution for the secondary structural elements in both lipid-free and cholesterol associated simulations (Figure 5A-B). We found that the α-helical propensity of the N-terminal region (residues 3-40, 48-90, 95-112, 118-124, 130-178) is highly conserved in both lipid-free and lipidated simulations.

**Figure 5:**
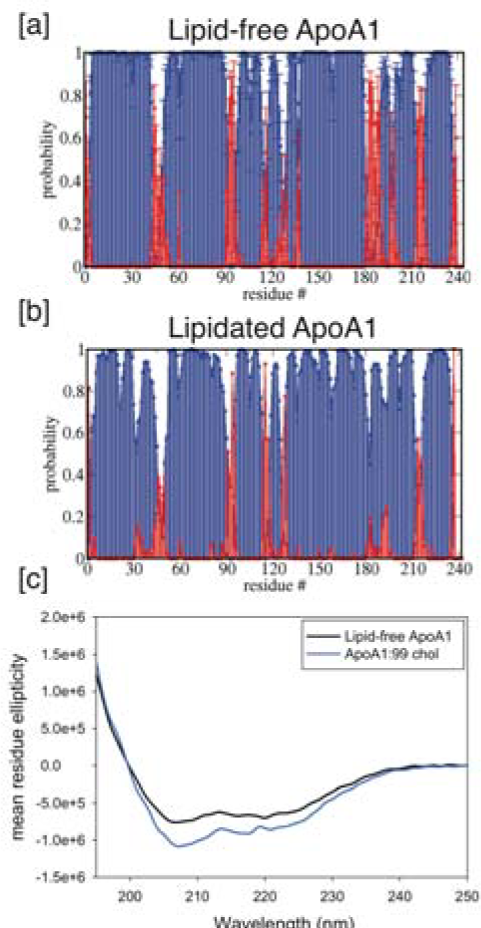
Monitoring change in secondary structure of ApoA1 upon interaction with cholesterol. A) Probability distribution for helix and coil secondary structure elements. The probability range is calculated on the basis of residue-based residence time, where 1 indicates the formation of helix (blue)/coil (red) secondary structure throughout the simulation. The error bars denote the standard deviation computed from five independent trajectories for the native structure’s simulations and B) Pilot lipidated simulation with ApoA1. (C) Mean residue molar ellipticity of monomeric ApoA1 with 99 molecules of cholesterol is plotted showing a drop at 208 nm and 222 nm, indicating increase in helicity upon cholesterol interaction.

The helical segments of ApoA1 lipid-free structure proposed in this study are largely in agreement with the secondary structure assigned based on previous experimental evidences (14-19, 26-31, 55-64, 70-77, 83-85, 93-97, 102-115, 147-148, 158-178, 224-227), as shown in Figure S11^23-27^. Interestingly, helical region 224-231, is also proposed to have the greatest lipid-binding property and contribution in initial lipid association^25, 27-29^. The region 115-150 was found to be sensitive to proteolysis and hence thought to be flexible and unstructured^30^. Previous reports also suggested that the N-terminal of the ApoA1 protein is relatively stable^31^. In contrast, C-terminal region was found to be highly unstructured in lipid-free, with residues 183-186, 189, and 216-217 found to be mostly in loop conformations. In particular, in the presence of cholesterol, residues (182-190, 198-199) dynamically interconvert between coil and helix (Figure S12). These findings largely agree with previous low resolution structural data where the C-terminal has been suggested to exist in a combination of helical and coil structure^15, 20, 31^.

Further, we also performed CD spectroscopy of lipid-free monomeric ApoA1 and, cholesterol bound ApoA1 (Figure 5C). The mean residue ellipticity drop at 208 nm and 222 nm, indicating increased α-helical region in the lipid-free monomeric state of ApoA1 compared to cholesterol bound ApoA1.

Overall, using combined experimental and simulation approach, we hereby showed that dynamics of ApoA1 protein domains is highly specific, indicating its link to distinct biological function. While the role of C-terminal is already established in previous reports^21-22^, our simulations reveal a novel binding site at the N-terminal. Furthermore, conformational map of structural transitions confirmed the existence of an obligatory intermediate state. Thus, deciphering the mechanisms of ApoA1 induced cholesterol recognition, a major component in RCT, will provide the means to significantly advance our understanding in maintaining cardiovascular health.

## Acknowledgements

We would like to thank Monika Verma for help in protein purification. LT is funded by INSPIRE Faculty grant from Department of Science and Technology under the LSBM-45 project. AR acknowledges the financial assistance from Council of Scientific and Industrial Research (CSIR), under the CARDIOMED-BSC0122 project. We are also thankful to CSIR-IGIB for infrastructure support and CSIR-4PI for supercomputing facility. LT also acknowledges adjunct faculty association with IIIT Delhi.

## Supporting Information Available

Description of the material included.

